# Mechanisms and heterogeneity of mineral use by natural colonies of the cyanobacterium *Trichodesmium*

**DOI:** 10.1101/2020.09.24.295147

**Authors:** Noelle A. Held, Kevin M. Sutherland, Eric A. Webb, Matthew R. McIlvin, Natalie R. Cohen, Alexander J. Devaux, David A. Hutchins, John B. Waterbury, Colleen M. Hansel, Mak A. Saito

**Affiliations:** MIT/WHOI Joint Program in Oceanography/Applied Ocean Science and Engineering, Woods Hole, MA 02543, USA; Department of Environmental Microbiology, Eawag, Dübendorf, Switzerland; Department of Environmental Systems Science, ETH Zürich, Zürich, Switzerland; Department of Earth and Planetary Sciences, Harvard University, Cambridge, MA 02138; Marine and Environmental Biology, Department of Biological Sciences, University of Southern California, Los Angeles, CA 90089, USA; Department of Marine Chemistry and Geochemistry, Woods Hole Oceanographic Institution, Woods Hole, MA 02543, USA; Department of Microbiology, Boston University School of Medicine, Boston, MA 02118, USA; Department of Biology, Woods Hole Oceanographic Institution, Woods Hole, MA 02543, USA

**Author notes:** Correspondence to: Mak Saito, **Email:**. **Author Contributions** N.A.H., M.M.M., J.W. and M.A.S. conceptualized the study. N.A.H. and M.M.M. performed proteomic analyses. K.M.S. and C.M.H. performed synchrotron-based analyses. N.C. performed trace metal and assisted with phosphate analyses. A.J.D. performed image analyses. E.A.W. and D.A.H. lead the Tricolim field expedition. N.A.H. and M.A.S. wrote the paper with input from all authors.

## Abstract

The keystone marine nitrogen fixer *Trichodesmium* thrives in high dust environments, and while experimental observations suggest that *Trichodesmium* colonies can access the essential nutrient iron from dust particles, it is not known the extent to which this occurs in the field. Here we demonstrate that *Trichodesmium* colonies actively process mineral particles in nature with direct molecular impacts. Microscopy and synchrotron-based imaging demonstrated heterogeneous associations with particles consistent with iron oxide and iron silicate minerals. Metaproteomic analysis of individual colonies revealed enrichment of biogeochemically-relevant proteins including photosynthesis proteins and metalloproteins containing iron, nickel, copper and zinc when particles were present. The iron-storage protein ferritin was particularly enriched implying accumulation of particle-derived iron, and multiple iron acquisition pathways including Fe(II), Fe(III), and Fe-siderophore transporters were engaged, including evidence of superoxide-driven particle dissolution. While the particles clearly provided iron, there was also evidence that the concentrated metals had toxic effects. The molecular mechanisms allowing *Trichodesmium* to interact with particulate minerals are fundamental to its success and global impact on nitrogen biogeochemistry, and may contribute to the leaching of particulate trace metals with implications for global iron and carbon cycling.

## Introduction

Marine nitrogen fixation is a key process that stimulates primary production in the N-depleted surface ocean, thereby influencing global carbon and nitrogen cycling^1,2^. First observed by mariners who referred to it as “sea sawdust,” the colonial cyanobacterium *Trichodesmium* is now recognized to be a major contributor to oceanic nitrogen fixation and therefore a crucial player in global nitrogen and carbon cycling^3,4^. Due to the high iron requirement of the nitrogenase enzyme, iron (Fe) is considered to be a dominant control on the distribution of nitrogen fixers, particularly *Trichodesmium*. However, mechanisms of iron uptake and utilization remain poorly understood^5–8^.

In nature, *Trichodesmium* forms large colonies that can reach several millimeters in size, with distinct morphologies including “puffs” and “tufts” (e.g. Fig. 1G, H). These colonies harbor complex microbiomes with diverse heterotrophic and phototrophic bacterial communities ^9,10^. The benefits of colony formation have been debated, but there is renewed interest in their utility for particulate iron acquisition^11–13^. This is important because *Trichodesmium* thrives in regions where Fe-rich continental dust is deposited such as the North Atlantic, Red Sea, and near land forms including Australia and Caribbean islands^14,15^. Furthermore, dust addition experiments have demonstrated that *Trichodesmium* puff colonies can acquire metals from iron (oxy/hydro)oxide particles, although the mechanistic underpinnings and biogeochemical importance of this process are not clear^8,11–13,16^.

**Figure. 1.**
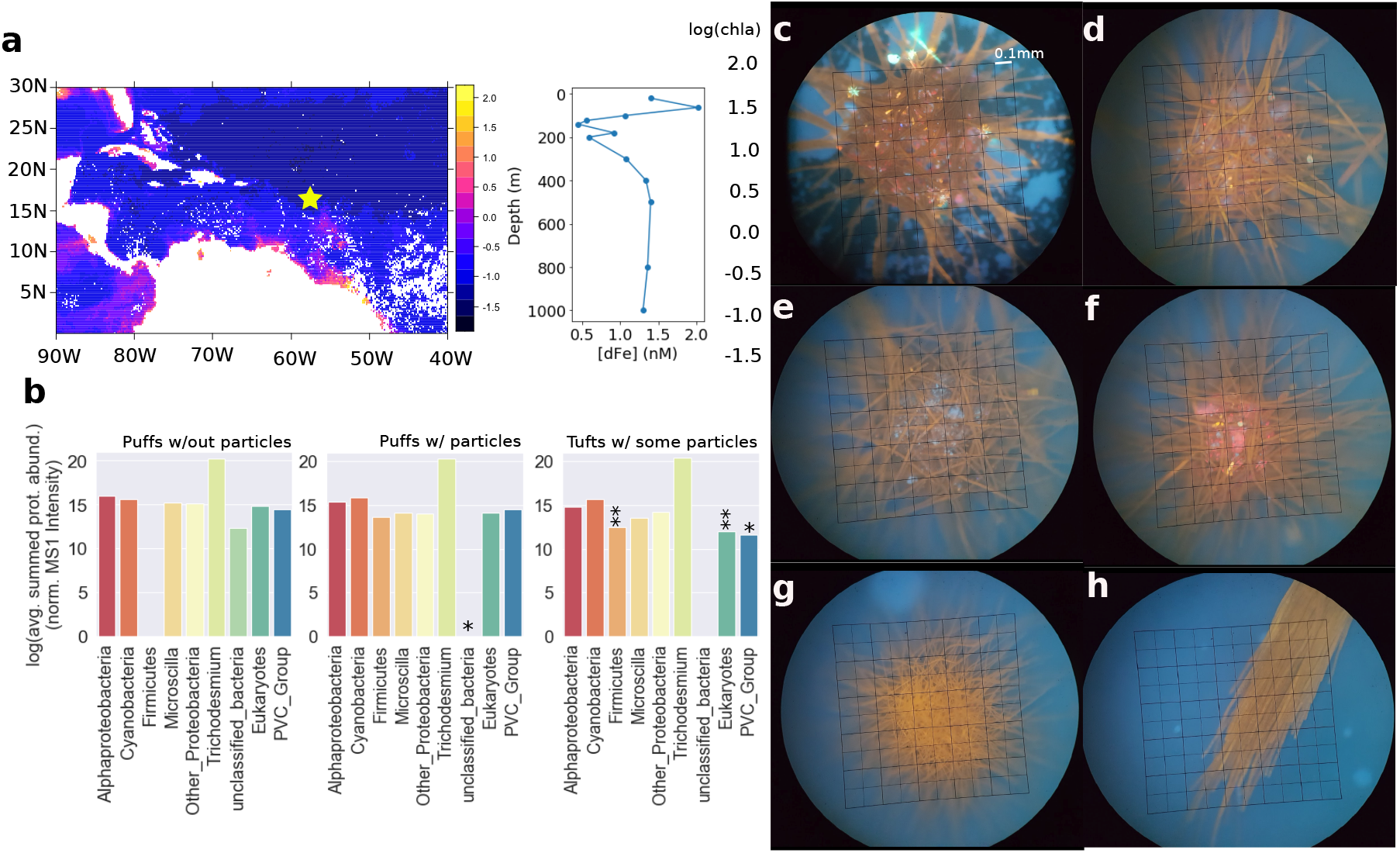
(A) Sampling location (yellow star) overlaid on MODIS-Aqua chlorophyll-A data averaged between March 1 and March 31, 2017, with dissolved iron profile at the sampling location. B) Biological diversity of the proteins identified in the single colony metaproteomes, with abundances of all detected proteins summed by major taxonomic groupings and separated by colony morphology. *indicates significant difference from puffs without particles at p < 0.1, ** the same at p < 0.05, Welch’s unequal variance t-test (C-F) Representative images of puff colonies with particles, (G) puff without particles and (H) a tuft with some particles. Images were collected in epifluorescent mode using a DAPI long pass filter set, without dyes. The scale bar in panel (C) applies to all images.

We investigated morphological and molecular differences in individual *Trichodesmium* colonies collected from a single plankton net conducted at 17:00 local time on March 11, 2018 in the tropical Atlantic Ocean in the vicinity of the Orinoco and Amazon river plumes (−57.5°W 16.5°N, Fig. 1A). Both puff and tuft colonies were present, though puffs dominated. The phosphate concentration was low (0.13 μM at 100m) as is typical in an oligotrophic environment, while the surface dissolved iron concentration was relatively high (2.02 nM at 100m), consistent with coastal or atmospheric inputs (Fig. 1A). The most abundant species at this location was an uncharacterized *Trichodesmium theibautii* species, as determined by *Trichodesmium-specific* metagenome-assembled-genome recruiting (see Table S2). Thirty individual colonies of mixed morphology were separated by hand-picking, immediately examined by fluorescent microscopy (385 excitation, > 420nm emission), then frozen individually for X-ray spectroscopy or metaproteomic analyses (Fig. S1). We observed that some colonies were associated with auto-fluorescent particles, which we hypothesized to be of mineral origin (Figure 1C-H). The particles fluoresced in the visual light range, appearing as yellow, red, or blue dots. Strikingly, colonies either had many such particles or none at all. In general the particles were concentrated in the center of puff type colonies, though they were also present in tufts but in in smaller numbers.

## Results and Discussion

Synchrotron based micro X-ray fluorescence (μ-XRF) element mapping of three representative particle-associated colonies demonstrated that the particles were enriched in iron (Fe), copper (Cu), zinc (Zn), titanium/barium (Ti/Ba, which cannot be distinguished by this method), and manganese (Mn), though the concentration approached the limit of detection for the latter element (Fig. 2). Iron concentrations were particularly high in the particles. Micro X-ray absorption near-edge structure (μ-XANES) spectra for Fe were collected on six particles - three each from two puffs (Fig. 2 and Fig. S2). The particles contained mineral bound iron with average oxidation states of 2.6, 2.7, two of oxidation state 2.9, and two of oxidation state 3.0 (Table S1). While the mineralogy of these particles could not be definitively resolved using μ-XANES, the structure of the absorption edge and post-edge region provided insight into broad mineral groups. Both Fe(III) (oxy/hydro)oxides and mixed-valence Fe-bearing minerals consistent with Fe silicates were present, suggesting heterogeneous mineral character. While we could not positively identify the silicate mineral phases based on XANES, the spectroscopic similarity of some samples to Fe-smectite and the geologic context suggest Fe-bearing clays were present (Fig. S4). In this geographic region, iron oxides could be sourced from atmospheric dust, and clays from the Orinoco and/or Amazon rivers. These colony-associated particles likely serve as a simultaneous source of nutritional (Fe, Ni, Co, Mn) and toxic (Cu) metals to the colonies. Release of metals from the particles likely vary over time, with Cu, Ni, Zn and Co continually leaching and Fe leaching initially, then re-adsorbing back onto particles unless organic chelates assist in solubilization^17^.

**Figure. 2.**
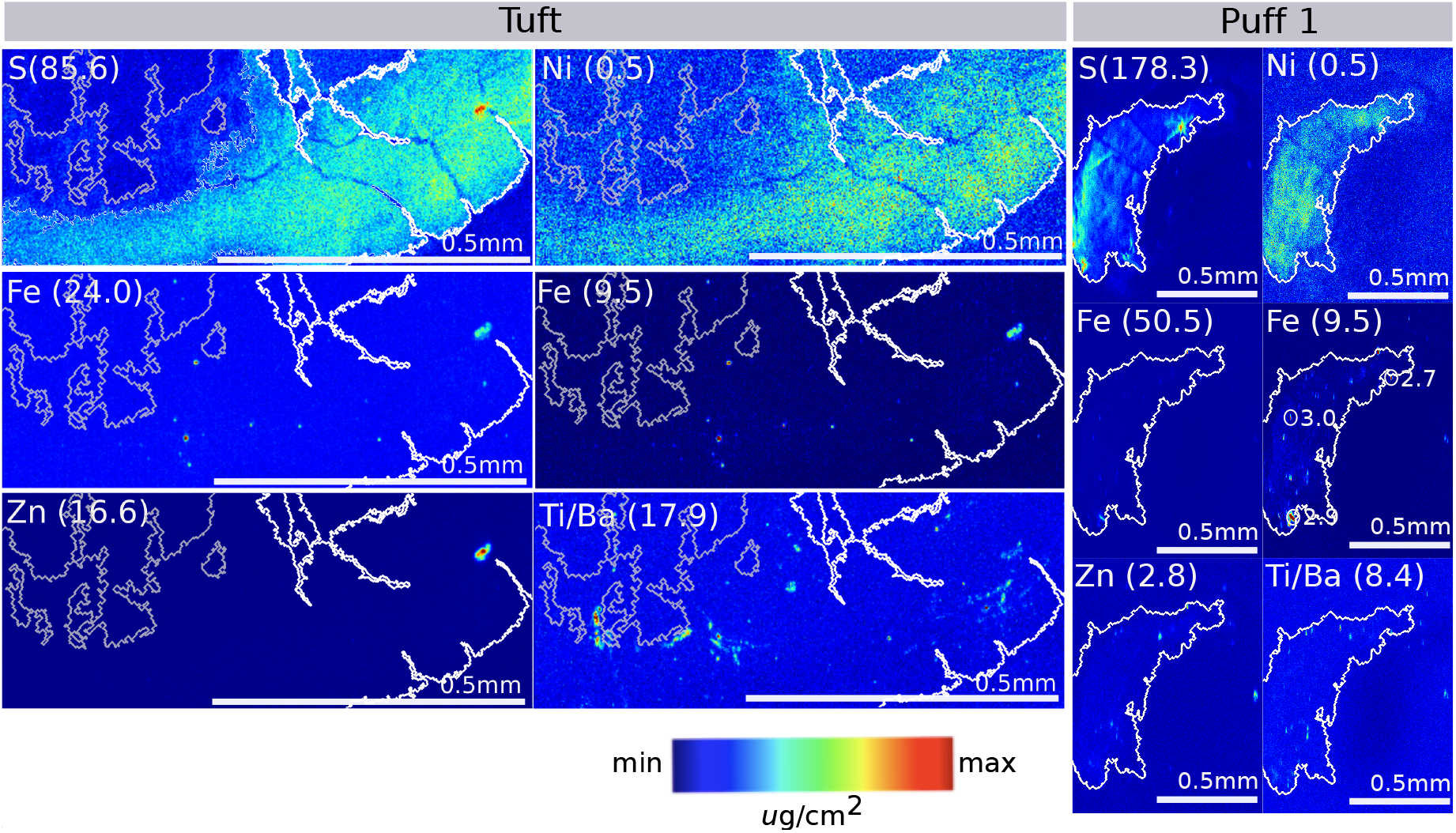
μ-XRF based element maps of a *Trichodesmium* tuft (left) and puff (right) colony (beamsize 3 x 3 mm). White/grey contours, based on the sulfur panel, which is indicative of biomass, have been provided (white = high [S] threshold, grey = lower [S] threshold). The color scale is the same for each image, with the maximum concentration for each element indicated in parentheses; iron is displayed using two scales. Iron oxidation states were determined via μ-XANES for three particles in the puff colony, and these are annotated. The corresponding XANES spectra are shown in Figure S4 and tabulated data in Table S1.

To understand the impact of the particles on colony diversity and function, we investigated metaproteomes of individual *Trichodesmium* colonies and their microbiota. Since all of the colonies were sampled from the same body of water, the presence of colonies with and without particles may be due to stochastics of the particle-colony encounter and/or sub-species differences among the colonies. Seven puffs without particles, fourteen puffs with particles, and four tufts with particles were analyzed by a new single colony metaproteomic method. Compared to population-level metaproteomes from this location, which each integrated 50-100 colonies, proteome depth for the low biomass single colonies was lower but sufficient for characterizing overall colony function (Fig. S6)^18^. In total, 1583 *Trichodesmium* and 487 epibiont proteins were identified across the 25 single colony metaproteomes versus 2944 *Trichodesmium* and 1534 epibiont proteins across triplicate population-level metaproteomes (Table S3 & S4). To ensure phylogenetic exclusivity, peptides used to identify epibiont proteins were ensured to be not present in the *Trichodesmium* genome (Table S5).

Together the single colony proteomes demonstrate a diverse and functionally active microbiome associated with the *Trichodesmium* colony (Fig. 1B and Fig. S7). Many commonly identified epibiont groups were present including Alphaproteobacteria, Microscilla, and non-*Trichodesmium* cyanobacteria. In general the epibiont community was similar among the colony types, however two clear differences emerged. First, firmicute proteins were more abundant in tufts and puffs with particles, suggesting enhanced, possibly anaerobic, metabolism. Second, Eukaryotic proteins were more abundant in puffs compared to tufts. These proteins likely represent copepods due to sequence similarity to the model organism *Calanus finmarchicus*, and this result is consistent with observed associations between copepods and puffs at this location (Fig. S7B). Notably, proteins from the PVC superphylum, which included Eukaryote pathogens such as an uncharacterized Chlamydia species, were also more abundant in puffs.

Particle presence was associated with clear differences in *Trichodesmium’s* proteome. In total, 131 proteins were differently abundant in puffs with versus without particles (p < 0.05, FDR-controlled Welch’s unequal variances t-test). Proteome differences were distributed across a variety of biogeochemically relevant proteins, including those related to four metals (Fe, Ni, Cu and Zn) and carbon fixation including the proteins rubisco (p = 0.04), citrate synthase (p = 0.01), and the accessory pigment allophycocyanin (p = 0.1). Particle presence did not affect nitrogenase abundance, consistent with laboratory dust addition experiments ^16^. There were indications that the particles altered nitrogen metabolism more generally, for instance through enrichment of nitrogen assimilation proteins glutamine synthetase (p = 0.008), spermidine synthase (p = 0.005), and a urea transporter (p = 0.02) (see Fig. S7)^18^. Because nitrogen fixation is a critical process, *Trichodesmium* may distribute iron to nitrogenase at a steady rate, while altering the activity of other systems in response to the particles.

The broadest and strongest response occurred in iron-related proteins, in agreement with the strong biogeochemical connection between iron and diazotrophy. Consistent with the high concentration of iron in the particles, several iron-containing proteins were significantly more abundant when particles were present, suggesting that the minerals were a micronutrient source. These included an iron-containing peroxidase (p = 0.03), the electron transport protein ferredoxin fdxH (p = 0.0006), and the iron storage/DNA binding protein Dps ferritin (p = 0.002) (Fig. 3). Increased abundance of ferredoxin is consistent with increased iron availability, since the non-iron containing flavodoxin typically substitutes during iron stress ^5,19^. Similarly, increased abundance of ferritin would serve to buffer and store iron acquired from the concentrated particulate metal source ^20^. Given the high iron demand of the nitrogenase metalloenzyme, the ability to store iron from rich but episodically available mineral particles could provide an important ecological niche in oceanic environments where iron can be scarce and its solubility is low.

**Figure 3.**
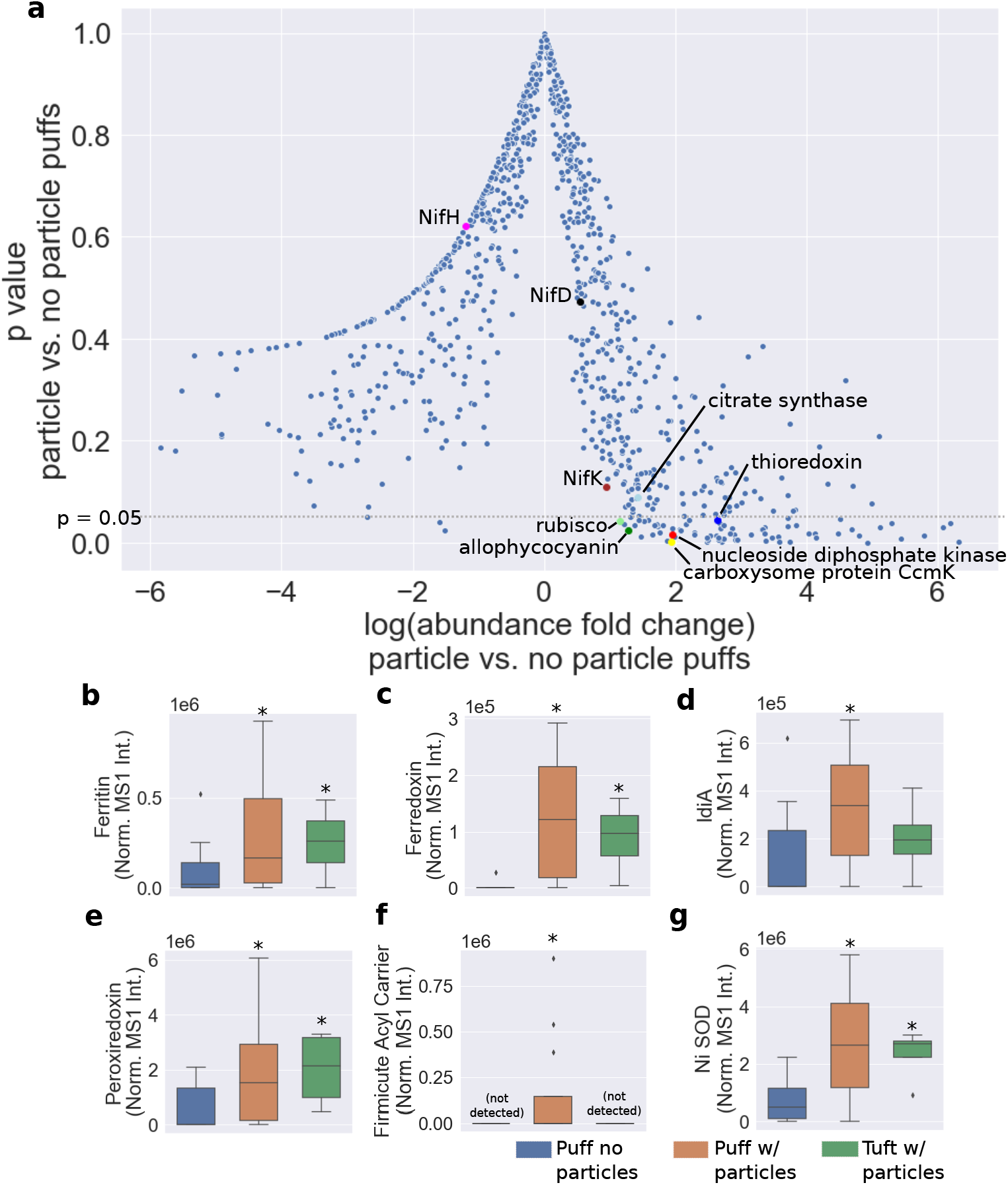
A) p value (Welch’s t-test) versus protein abundance reported as fold change for puffs with vs. without particles. Below the grey dotted line (p = 0.05), the differences are statistically significant. Positive fold change indicates the protein was more abundant when particles were present. Proteins of interest are highlighted. (B-E) Relative abundance of selected proteins for the different colony types, presented as box plots (center line = median, box limits = first and third quartiles, whiskers = data min and max, diamonds = outliers). *Indicates statistically significant difference compared to the puffs without particles, p < 0.05.

Multiple uptake mechanisms were involved in obtaining iron from the mineral particles. Iron acquisition in *Trichodesmium* is not well understood but at least three systems are known: the FeoB system (Fe(II)), the IdiA system (Fe(III)), and uptake via Fe-siderophores ^8,21^. The single colony metaproteomes provided evidence for the latter two mechanisms; FeoB is rarely identified in metaproteomes of diazotrophs, possibly due to its low copy number/high efficiency (Fig. 4)^19,22,23^. Despite evidence that the particles provided iron to the colonies, the transport protein IdiA was more abundant during particle associations (Fig. 3D). IdiA is often used as a biomarker of iron stress because it is responsive to/more abundant in low Fe conditions^18,19,23^. Due to its specificity for Fe(III), IdiA is likely involved in direct acquisition of iron from the particles. Additionally, iron pulses from the particles may have been recent, and IdiA may not yet have been turned over or diluted. There was also evidence that iron-binding siderophore systems derived from the colony microbiome were involved: a Firmicute acyl carrier protein putatively involved in siderophore production was enriched in puffs with particles (Fig. 3F, p = 0.04), and a TonB dependent transporter (TBDT) for ferrienterochelin/colicins was identified only in puffs with particles^24^. This corroborates earlier evidence that though *Trichodesmium* does not produce its own siderophores, it acquires siderophore-bound iron produced via mutualistic interactions with epibionts, especially when provided with concentrated dust^8,25^.

**Figure 4.**
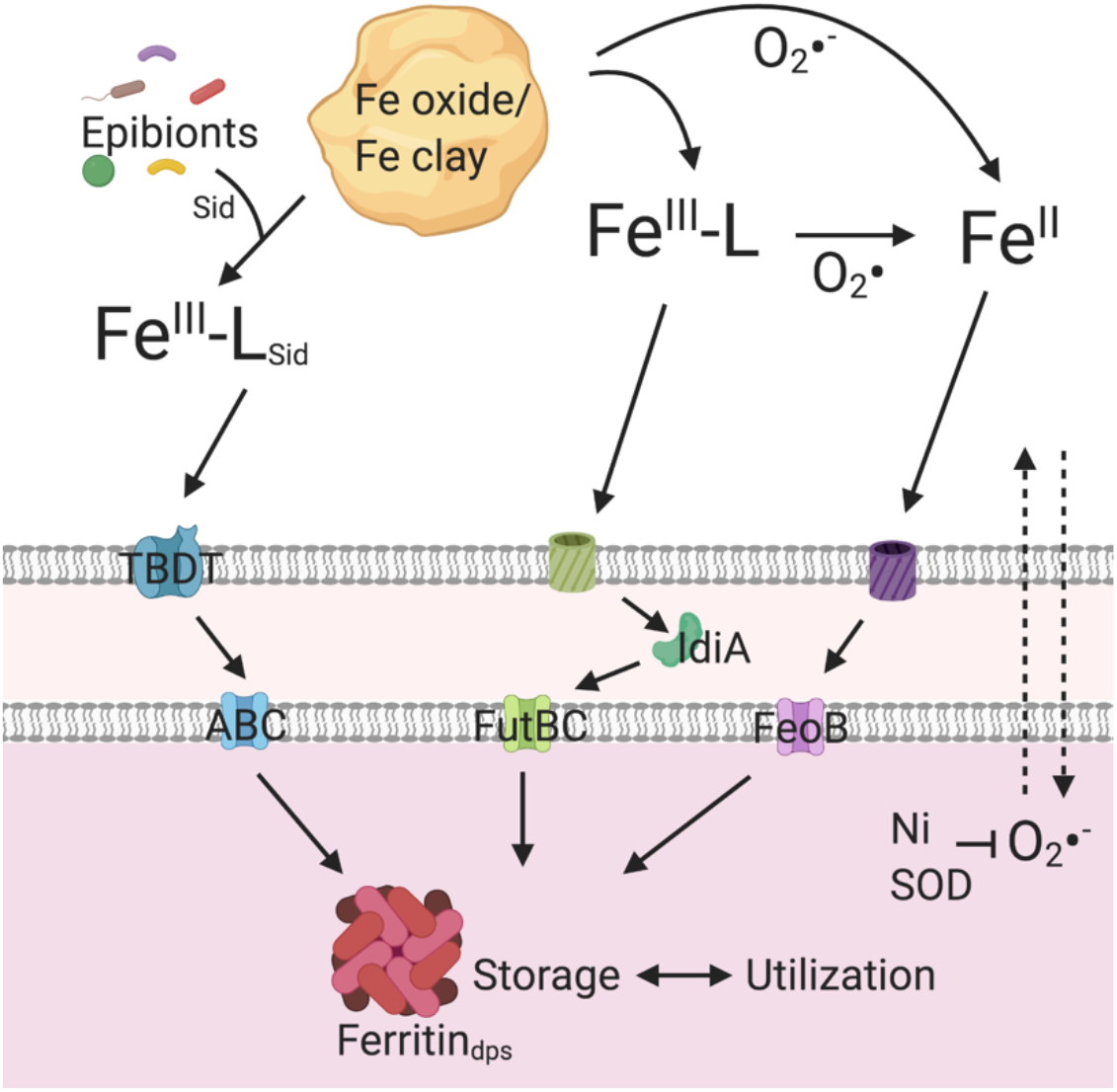
Model of iron uptake mechanisms involved in utilization of particle-derived iron. Fe(III)-L and Fe(II) are thought to enter the periplasm through passive porins/receptors. ^8^ Proteins with annotated names are identified, otherwise the following general functional names are used: TBDT = unspecified TonB dependent transporter, ABC = unspecified ABC transporter, Ni SOD = nickel superoxide dismutase, which responds to increased superoxide production during high productivity/particle dissolution.

Nickel superoxide dismutase (Ni SOD) was significantly more abundant in particle-associated colonies, reflecting the need to regulate the reactive oxygen species (ROS) superoxide during particle dissolution/enhanced productivity (Fig. 3G p = 0.004). There are multiple reasons for elevated superoxide production in particle-associated colonies. First, and most directly, *Trichodesmium* produces extracellular superoxide to enhance particle dissolution/reduction of Fe(III)^8^. Second, enhanced productivity, as was suggested based on enrichment of photosynthesis proteins, is associated with growth-related extracellular superoxide production^8^. In addition to extracellular ROS, *Trichodesmium* has high intracellular ROS during photosynthesis due to the fact that it has among the highest Mehler reaction activity of any photosynthetic organism^26^. Puffs in particular are known to closely regulate superoxide production with a possible link to cell signaling and growth^27^. Further supporting a redox-centered explanation, other regulators including peroxiredoxin and thioredoxin were more abundant in puffs with particles (see Fig. 3E, p = 0.03 and 0.04, respectively). Additionally, the increase in Ni SOD may reflect that the particles were providing nickel, an essential nutrient that can be limiting to *Trichodesmium*^28^.

Together, the molecular evidence points to a model in which *Trichodesmium* colonies differentiate between dissolved versus particulate iron. First, consistent with the mineralogical profile of the particles, the colonies engaged specific iron uptake mechanisms that prioritized the Fe(III) state as well as balancing cellular redox status during bio-enhanced Fe(III) reduction (Figure 4). Then, once inside the cell, mineral-derived iron was preferentially stored via ferritin. This finding adds complexity to the canonical regulatory model that IdiA and ferritin exist on a continuum with IdiA abundant during iron limitation and ferritin present only when iron is replete. It suggests that *Trichodesmium* alters uptake and utilization mechanisms in response to the iron’s coordination environment. Whether or not this distinction is made directly, i.e via a specific mineral sensing mechanism, or indirectly due to increased intracellular metal concentration, is not yet known. Either way, it is clear that multiple metabolic and homeostasis systems were involved in this coordinated response to the presence of particulate iron.

Other intriguing aspects of particle associated *Trichodesmium* colonies included the deployment of Cu and motility related proteins. *Trichodesmium* is extraordinarily sensitive to Cu toxicity, and close proximity to mineral particles was associated with enrichment of the copper chaperone/homeostasis protein CopZ (Fig. S9D)^29,30^. Evidence also pointed to involvement of putative movement proteins including RTX proteins and the chemotaxis regulator CheY, both enriched in puffs with particles (Fig. S9E-H). This supports recent observations of *Trichodesmium* cilia transporting particles towards the puff center, indicating that this behavior may be induced by motility two-component regulatory systems, many of which have unknown environmental cues.

This study provides direct evidence that *Trichodesmium* colonies actively capture and process oxides and silicates under natural conditions, and that key aspects of the colony’s proteome respond as a result. *Trichodesmium* typically thrives in high dust environments and this study provides a biochemical basis for this specialized niche. The results have geochemical implications beyond *Trichodesmium* biology. Specifically, active capture and degradation of mineral particles may increase iron availability in the oligotrophic surface ocean. In this way, abundant *Trichodesmium* colonies may have an important role in the leaching of particulate trace metals with implications for global iron and carbon cycling^31^. The molecular scale single colony analyses presented here present a more sophisticated perspective of *Trichodesmium’s* response to mineral particles that could be leveraged in future biogeochemical models. Particulate iron utilization by *Trichodesmium* appears to be a critical niche, and is likely a significant factor determining this organism’s ecological success and fixed nitrogen contributions to the global ocean.

## Materials and Methods

### Sampling and microscopy

All of the colonies used in this study were sampled from a single plankton net conducted at −57.5°W 16.5°N at 17:00 local time on March 11, 2018 on the AT39-05/Tricolim expedition (R.V. Atlantis, Chief Scientist D. Hutchins, https://www.bco-dmo.org/deployment/765978). A 130 μm net was released to approximately 20 m depth, then pulled back to surface and the process repeated five times. Colonies were hand picked by gentle pipetting, rinsed twice in 0.2 μm-filtered trace metal seawater, and decanted into 0.2 μm sterile filtered trace metal seawater until imaging. All at-sea colony picking and handling was conducted in a Class 100 trace metal clean environment. Colonies were imaged with a Zeiss epifluorescent microscope using transmitted light and/or a long-pass fluorescent filter set. At the time of imaging they were labeled as “particle containing” or not and classified as puffs or tufts. They were then decanted individually onto trace metal clean 0.2μm Supor filters and flash frozen in liquid nitrogen, then stored at −80°C until analysis at the home laboratory. Images were captured with a Samsung Galaxy Note 4 using a SnapZoom universal digiscoping adapter. MODIS-Aqua data for Fig. 1A was obtained from the NASA Goddard Space Flight Center, Ocean Ecology Laboratory, Ocean Biology Processing Group; (2014): Sea-viewing Wide Field-of-view Sensor (SeaWiFS) Ocean Color Data, NASA OB.DAAC Accessed on 2020-04-08.

### Nutrient and trace metal analyses

Seawater was sampled using a trace metal clean rosette consisting of Niskin-X bottles. Niskins were pressurized with nitrogen gas in a shipboard Class 100 clean room and seawater was filtered through 0.2 μm Supor membranes to remove particles. Aliquots for macronutrient (phosphate) analysis were frozen immediately at sea and were thawed just prior to analysis. Phosphate was quantified using a Technicon Autoanalyzer II by Joe Jennings at Oregon State University. Aliquots for dissolved metal analysis were acidified with concentrated trace metal clean HCl (Seastar) to pH 1.8 and allowed to equilibrate for ~1 month prior to analysis. Dissolved iron was concentrated using a seaFAST automated pre-concentration system and quantified on an ICAP Q inductively coupled plasma mass spectrometer (ICP-MS).

### Sample handling and preparation for single colony proteomics

Upon return to the lab, the colonies were carefully cut out of the filter to reduce the volume of liquid needed for protein extraction. The filter sections were submerged in PBS buffer with 10% sodium dodecyl sulfate (SDS), 1 mM magnesium chloride, 2 M urea and benzonase nuclease, heated at 95 °C for 10 min, then shaken at room temperature for one hour. Proteins in the resulting supernatant were quantified by the BCA assay. The proteins were digested with a modified tube gel protocol following Saito et al., 2014, but instead of the typical 200 μL final volume only 50 μL final volume was used.^32,33^ Additionally, the protein precipitation/purification step was eliminated because this is another source of total protein loss. Instead, the samples were treated with benzonase nuclease during the initial extraction phase to solubilize any DNA/RNA components, allowing the purification step to be skipped. Briefly, the proteins were embedded in an acrylamide gel, washed with a 50:50 acetonitrile: 25 mM ammonium bicarbonate buffer, dehydrated by acetonitrile treatment, then treated for one hour at 56 °C with 10 mM dithiothreitol in 25 mM ammonium bicarbonate followed by one hour at room temperature with 55 mM iodacetamide. Gels were dehydrated again and rehydrated in trypsin (Promega Gold) at a ratio of 1 μg trypsin: 20 μg total protein in 25 mM ammonium bicarbonate. Proteins were digested overnight at 37 °C with shaking. The peptides were then extracted from the gels in 20 μL peptide extraction buffer (50% acetonitrile, 5% formic acid in water). The resulting peptide mixtures were concentrated to 0.2 μg total protein/μL final concentration. While the tube gel method was used for samples presented here, magnetic bead and soluble protein digestion methods were also tested. Total protein recovery was lower with these methodologies, perhaps because these methods do not use SDS, which in our hands is a good lysing agent for *Trichodesmium*.

### LC-MS/MS analysis

Metaproteome analyses were conducted by tandem mass spectrometry on a Thermo Orbitrap Fusion using 0.5 μg total protein injections and a one-dimensional 120 min non-linear gradient on a 15 cm C18 column (100 □m x 150 mm, 3 μm particle size, 120 Å pore size, C18 Reprosil-Gold, Dr. Maisch GmbH packed in a New Objective PicoFrit column). LC lines were shortened when possible to reduce the possibility of sample loss to the tubes. Blanks were run between each sample to avoid carryover effects. For each run 0.5 μg of protein was injected directly the column using a Thermo Dionex Ultimate3000 RSLCnano system (Waltham, MA); if less than 0.5 μg of protein was available, the entire sample was injected. The samples were analyzed on a Thermo Orbitrap Fusion mass spectrometer with a Thermo Flex ion source (Waltham, MA). MS1 scans were monitored between 380-1580 m/z, with a 1.6 m/z MS2 isolation window (CID mode), 50 millisecond maximum injection time and 5 second dynamic exclusion time. The resulting spectra have been deposited to the ProteomeXchange Consortium via the PRIDE partner repository with the dataset identifier PXD016330 and 10.6019/PXD016330.^34^

### Bioinformatics analyses

The spectra were searched using the SEQUEST algorithm with a trimmed *Trichodesmium* sequence database. To generate the sequence database, triplicate Tricho-enriched metaproteomes from the same location (aka “population biomass” samples), each integrating ~50-100 colonies hand-picked from the same plankton net, were analyzed using a publicly available *Trichodesmium* consortia metagenome collected at Station BATS (IMG ID 2821474806). Based on these population metaproteomes, the sequence database was trimmed to include only the proteins identified at a 1% protein and peptide FDR level, with protein scoring calculated in Scaffold (Proteome Software, Inc). Hand-refined metagenome-assembled genomes (MAGs) from *Trichodesmium* populations throughout the AT39-05 transect were also included in the search: these included four *Trichodesmium theibautii* species (one H94 species and three uncharacterized *T. theibautiis*) and 17 MAGs from the epibiont community. The single colony metaproteomes were searched using the SEQUEST search engine with parent mass tolerance +/-10ppm and fragment mass tolerance 0.8 Dalton, allowing cysteine modification of +57.022 Daltons and methionine modification of +16 Daltons. The results were statistically validated at the 1% FDR level using the Scaffold program. This resulted in 1495 protein identifications across the individual colonies. When the whole *Trichodesmium* consortia metagenome was used, only 800 proteins were identified at the 1% protein and peptide FDR level, so reducing the search space significantly improved data quality.^35,36^ When the Trichodesmium MAGS were also included the single colony metproteomes resulted in 2075 proteins identified. Peptides used to identify *Trichodesmium* proteins were determined to be phylogenetically exclusive to the genus using the open source Metatryp software package last accessed on May 23, 2020.^37^ Statistical tests (Welch’s t tests) were performed using the Scipy stats python library and the results are reported in Table S2. P-values were FDR controlled by the Benjamini-Hochberg procedure; at alpha = 0.05 and alpha = 0.1, the calculated FDR was 0%; the FDR rose to 0.56% by alpha = 0.25.

### Micro-X-ray fluorescence and Micro-X-ray absorption spectroscopy

Micro-X-ray fluorescence (μ-XRF) and micro-X-ray absorption spectroscopy (μ-XAS) were conducted at the Stanford Synchrotron Radiation Lightsource (SSRL) on beamline 2-3 with a 3 μm raster and a 50 ms dwell time on each pixel. μ-XRF data were analyzed using MicroAnalysis Toolkit.^38^ Elemental concentrations were determined using standard foils containing each element of interest. The relative proportions of Fe(II) and Fe(III) were determined by fitting the edge position of the background subtracted, normalized XANES spectra. Fe XANES spectra were fit using the SIXPACK Software package^39^, and redox state was estimated by conducting a linear combination fitting of the absorption edge (7115-7140 eV) using the model compounds siderite (FeCO_3_) and 2-line ferrihydrite as end-member representatives of Fe(II) and Fe(III), respectively. Further, these values were confirmed through deconvolution of the edge shape using Gaussian peaks at two fixed energies corresponding to primary Fe(II) (7122 eV) and Fe(III) (7126 eV) contributions (PeakFit software, SeaSolve Inc.).^40^ Linear combinations of the empirical model spectra were optimized where the only adjustable parameters were the fractions of each model compound contributing to the fit. The goodness of fit was established by minimization of the R-factor.^41,42^

Although mineral identity cannot be conclusively determined with XANES, visual comparison of the edge features are indicative of broad Fe-bearing mineral groups including many common oxide and silicate minerals. Thus, to get a general sense of mineral groups, we have included several of the top spectral library query hits (Figure S4) for the six particles we looked at in this study. These include 2-line ferrihydrite and goethite (as Fe oxyhydroxide phases), ferrosmectite (as Fe-bearing secondary clays), and biotite (as a primary silicate). We also included siderite (as an Fe(II)-bearing carbonate) in Figure S4 for comparison to a pure Fe(II) phase.

### Image contour analysis for element concentration maps

Image contours were generated for the sulfur element concentration maps in Figure 2 using the Scikit-image python library^43^. First, using the contrast histograms for the sulfur images, algorithmically-defined thresholds were applied. For the tuft image in Fig. 2, two thresholds were used to capture the high and low biomass regions. The resulting binary images were then morphologically dilated to remove noise and connect gaps between objects. A list of contours for the binary image was generated using the marching squares/cubes algorithm^44^. The longest contours were overlaid on the element maps from which they were generated.

## Supporting information

Supplemental Discussion, Figures, Table S1

## Data availability

All data are provided in the main text or as supplementary materials. Additionally, the mass spectrometry proteomics data have been deposited to the ProteomeXchange Consortium via the PRIDE partner repository with the dataset identifier PXD016330 and 10.6019/PXD016330. The processed proteomics data can also be accessed at BCO-DMO (DOI: 10.26008/1912/bco-dmo.786694.1).

## Code availability

Data analyses and plotting were conducted in Python 3.0 (https://www.python.org/) using the pandas (https://pandas.pydata.org/), matplotlib (https://matplotlib.org/), seaborn (https://seaborn.pydata.org/),scipy stats (https://scipy.org/), and scikit image (https://scikit-image.org/) libraries. Fully reproducible example code, including statistical analyses, can be found at https://github.com/naheld/Tricho_Singlecolony_MetaP.

## Acknowledgments

We thank the science team and crew of the Tricolim cruise, especially Jaclyn Saunders, Michael Mazzotta, and Asa Conover. Joe Jennings (OSU) performed the macronutrient analysis. This work was supported by NSF Graduate Research Fellowship grant # 1122274 [N.Held], the Gordon and Betty Moore Foundation (grant number 3782 [M.Saito]) the National Science Foundation (grant numbers OCE 1657766 [M.Saito] and OCE 1657757 [D.Hutchins, E.Webb]), the Woods Hole Oceanographic Institution Ocean Ventures Fund [N. Held] and the Simons Foundation (grant 544236 to N. Cohen). Use of the Stanford Synchrotron Radiation Lightsource, SLAC National Accelerator Laboratory, is supported by the U.S. Department of Energy, Office of Science, Office of Basic Energy Sciences under Contract No. DE-AC02-76SF00515.

