## Supplemental Discussion, Figures, Table S1 for "Mechanisms and heterogeneity of mineral use by natural colonies of the cyanobacterium *Trichodesmium*"

### Supplementary Information for Held et al. "Mechanisms and heterogeneity of mineral use by natural colonies of the cyanobacterium *Trichodesmium*"

#### Supplementary Discussion

##### Taxonomic attributions of eukaryotic proteins from the epibiont community

For some of the eukaryote epibiont proteins, taxonomic attributions appeared to be non-marine. However, the proteins are more likely related to zooplankton not present in the genomic database since they included conserved actin, tubulin, ATP synthase, and histone proteins. Many of the proteins have homologs in the copepod model *Calanus finmarchicus,* such as histone protein TCCM_0779.00001180 (BLAST hit to *C. finmarchus* histone protein, E = $1e^{-4}$) and tubulin protein TCCM_0148.00002430 (BLAST hit to *C. finmarchus* alpha-tubulin, E = $4e^{-23}$). Copepods are phylogenetically diverse, and few example genomes exist, explaining the lack of coverage in metagenome annotations.[1] Even after seawater rinsing, we observed many colony associated copepods, particularly on puff-type colonies (see Fig. S7B). Copepods are known predators of *Trichodesmium*; certain species have specialized hooks for grabbing filaments and one species, *Macrosetella gracilis,* houses its eggs in *Trichodesmium* filaments.[2,3]

##### Possible involvement of motility proteins in entraining mineral particles

Recent laboratory studies have visually observed colonies to shuttle particles to the colony center on the order of minutes, and that this involves complex rotation, bending, stretching, and flipping movements of trichomes which are in constant motion within the colonies.[4,5] While these fascinating observations implied a specific response to particles by *Trichodesmium*, the mechanistic basis for this behavior was not known. Here, we observed a possible effect of particle presence on the abundance of the twitching motility response regulator PilH, which may regulate the direction of the pilus motor. PilH was abundant only in puffs with particles, and not identified at all in puffs without particles or in tufts (Fig. S9). This implied that *Trichodesmium's* motility machinery may be actively involved in particle entrainment, either by translocating particles along the trichome or via collective movement of the trichomes to push the particles to the colony center. While *Trichodesmium* has a complete set of genes encoding type IV pili, only two other pili proteins were observed - PilT pili retraction protein and PilM pili assembly protein. These structural proteins were not significantly more or less abundant when particles were present, implying that particle entrainment may occur simply through PilH regulation of pilus motor movement (Fig. S10). In general, *Trichodesmium's* motility is not well understood; for instance, despite having chemotaxis sensory tranducers, *Trichodesmium* lacks the flagellin structural protein with which they interact. It also contains 25 putative RTX proteins that are related to the SwmA motility protein found in marine *Synechococcus,* one of which was observed and was highly abundant when particles were present (Fig. S9).[6,7]

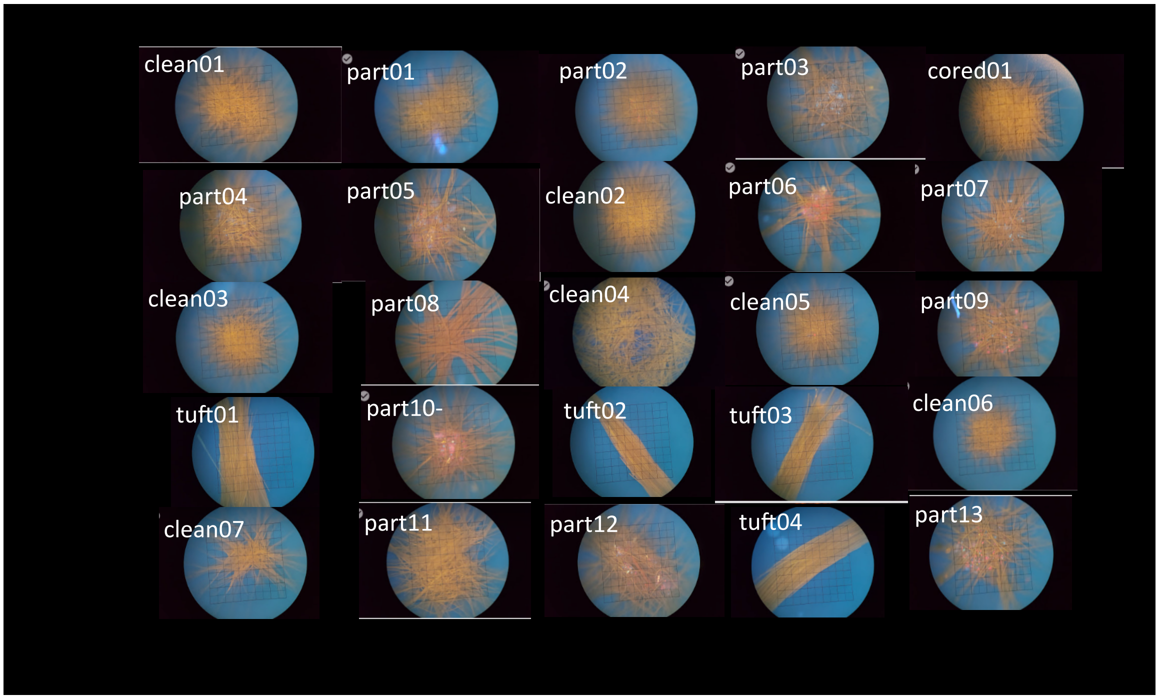


**Figure S1.** DAPI long pass filter images for all of the colonies examined in this study.

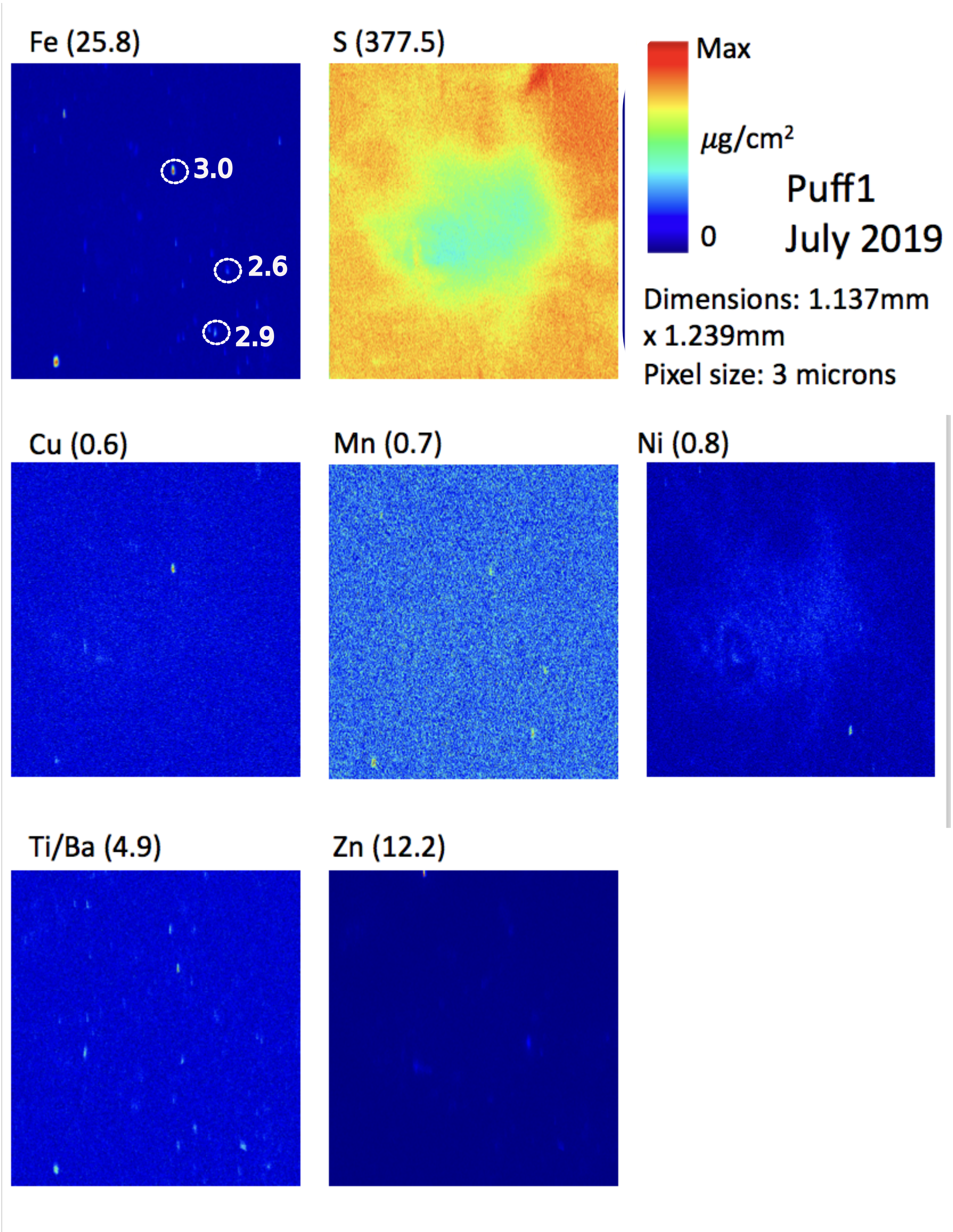


**Figure S2.** µ-XRF element mapping of a second *Trichodesmium* puff colony. Iron oxidation states were determined for three particles and are annotated in the respective panel. The corresponding XANES spectra are shown in Figure S5 and tabulated data in Table S1. Note that biomass contours are not provided for this puff, because as reflected in the sulfur panel and in figure S5, the entire mapped area contains *Trichodesmium* biomass.

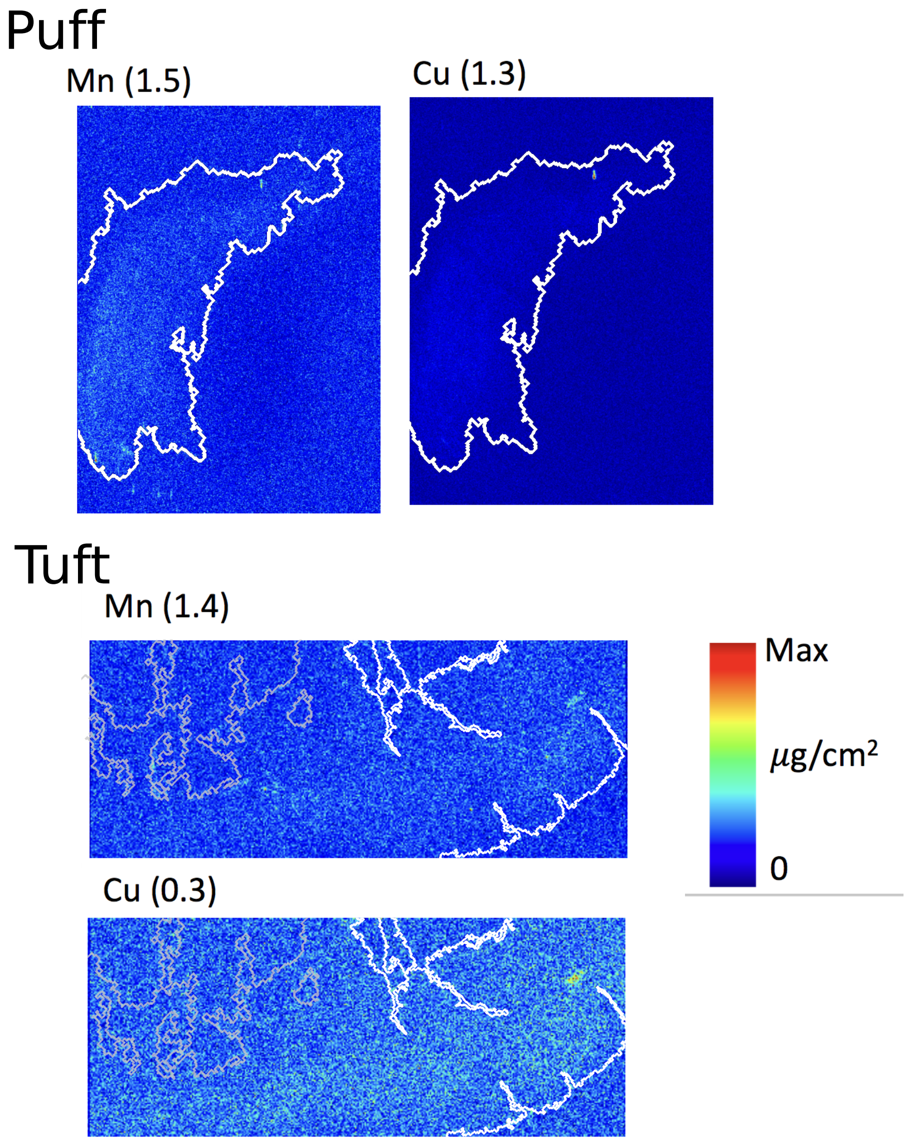


**Figure S3.** Additional µ-XRF element maps of the *Trichodesmium* tuft and puff colony featured in Figures 8 and 9, respectively. The maximum of the color scale is given next to each element. Contours have been drawn as in Fig. 2.

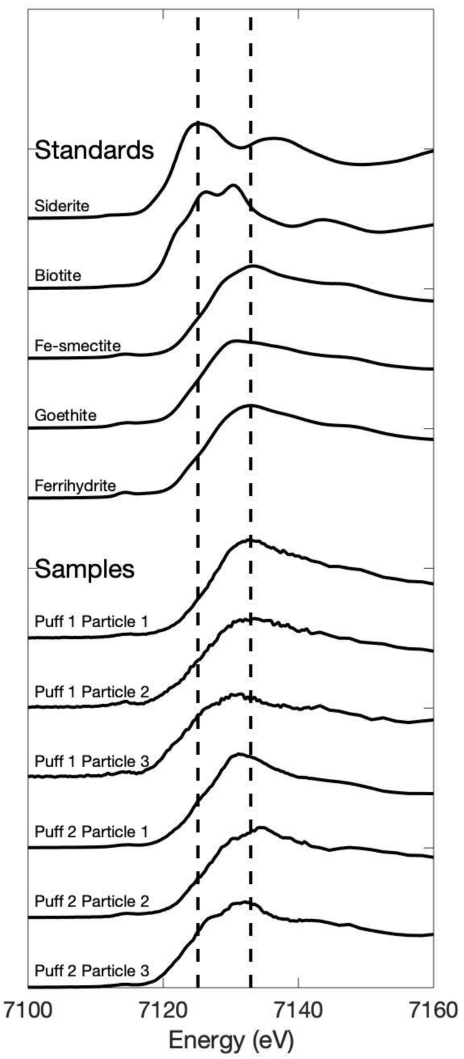


**Figure S4.** Fe K-edge μ-XANES spectra of the particles in Figures 2 and S3 compared to model standard spectra. The dotted vertical lines represent peak energies for Fe(II) and Fe(III). The doublet peak feature accentuated in biotite is also seen in smectite, and clearly visible in some of the puff particle spectra, such as puff 1 particle 3, and puff 2 particles 2 and 3. In contrast, oxides are dominated by a clean white line and single peak as observed for ferrihydrite and for instance puff 1 particle 1.

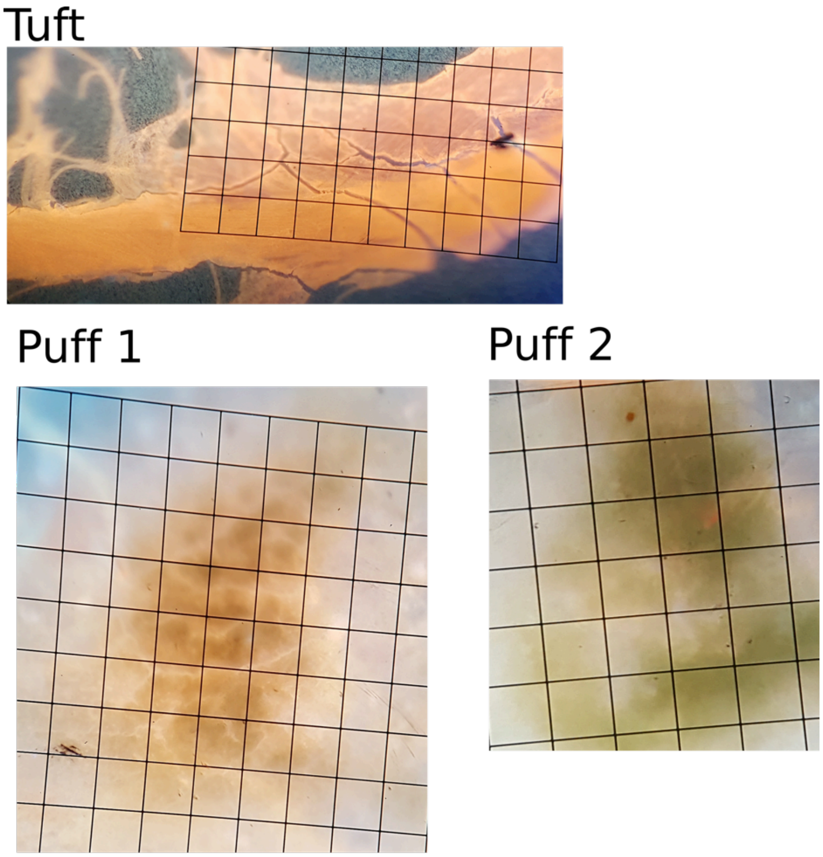


**Figure S5.** Fluorescent microscopy images of the tuft and two puffs used for the SSRD analyses. These images were captured after the colonies were freeze-dried using a DAPI long pass filter but no dyes. Note that the cells lysed during the freeze-drying process.

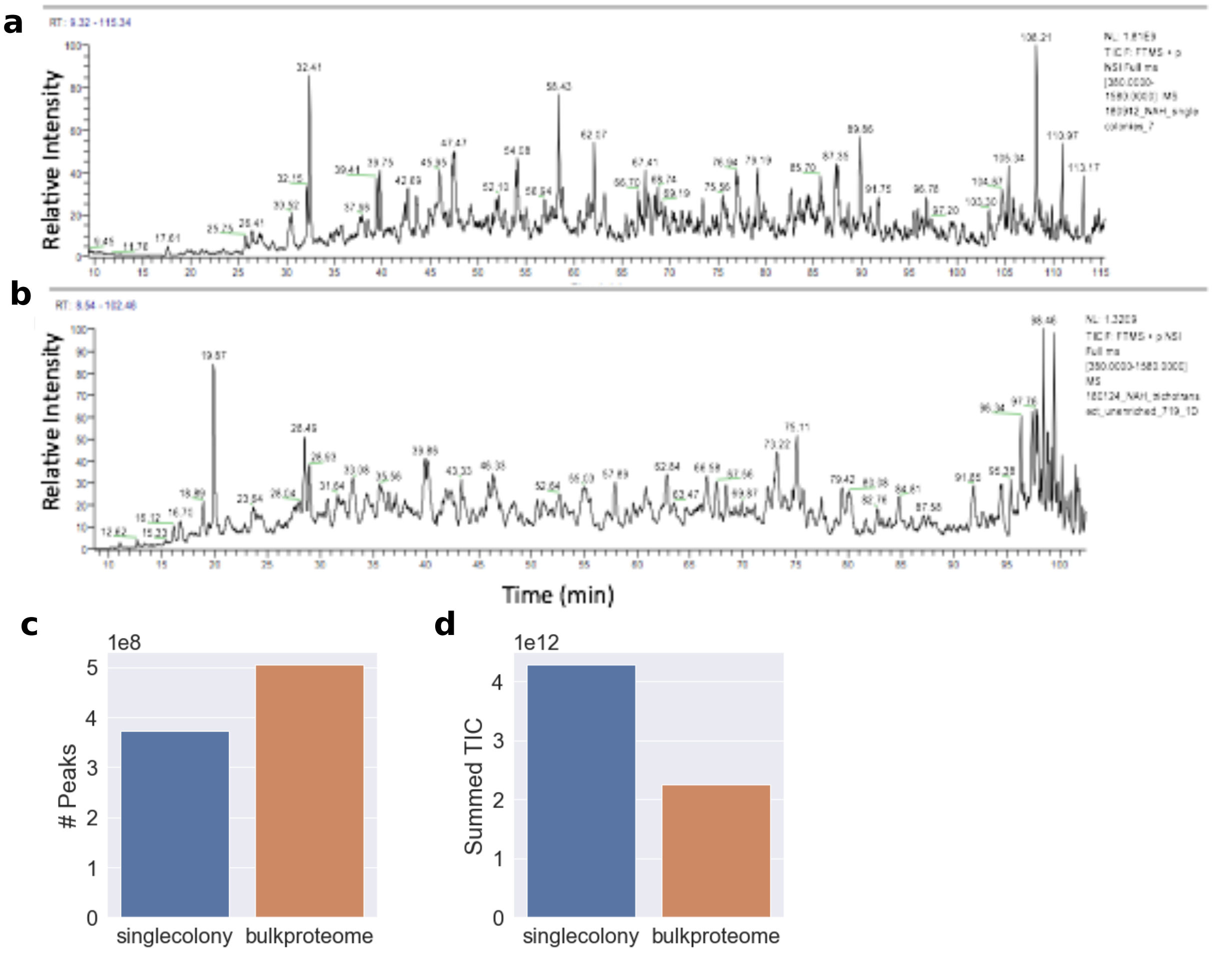


**Figure S6.** Chromatographic traces for (A) a single colony and (B) bulk *Trichodesmium* metaproteome from the same station. (C) number of peaks identified in the spectrum and (D) summed total ion current (TIC) for the two samples. Peak counting was performed following Saito et al., 2018.9 The single colony metaproteome had higher total intensity and slightly fewer peaks identified. The sample was complex, illustrating the challenge in acquiring high quality data from a small amount of field sample.

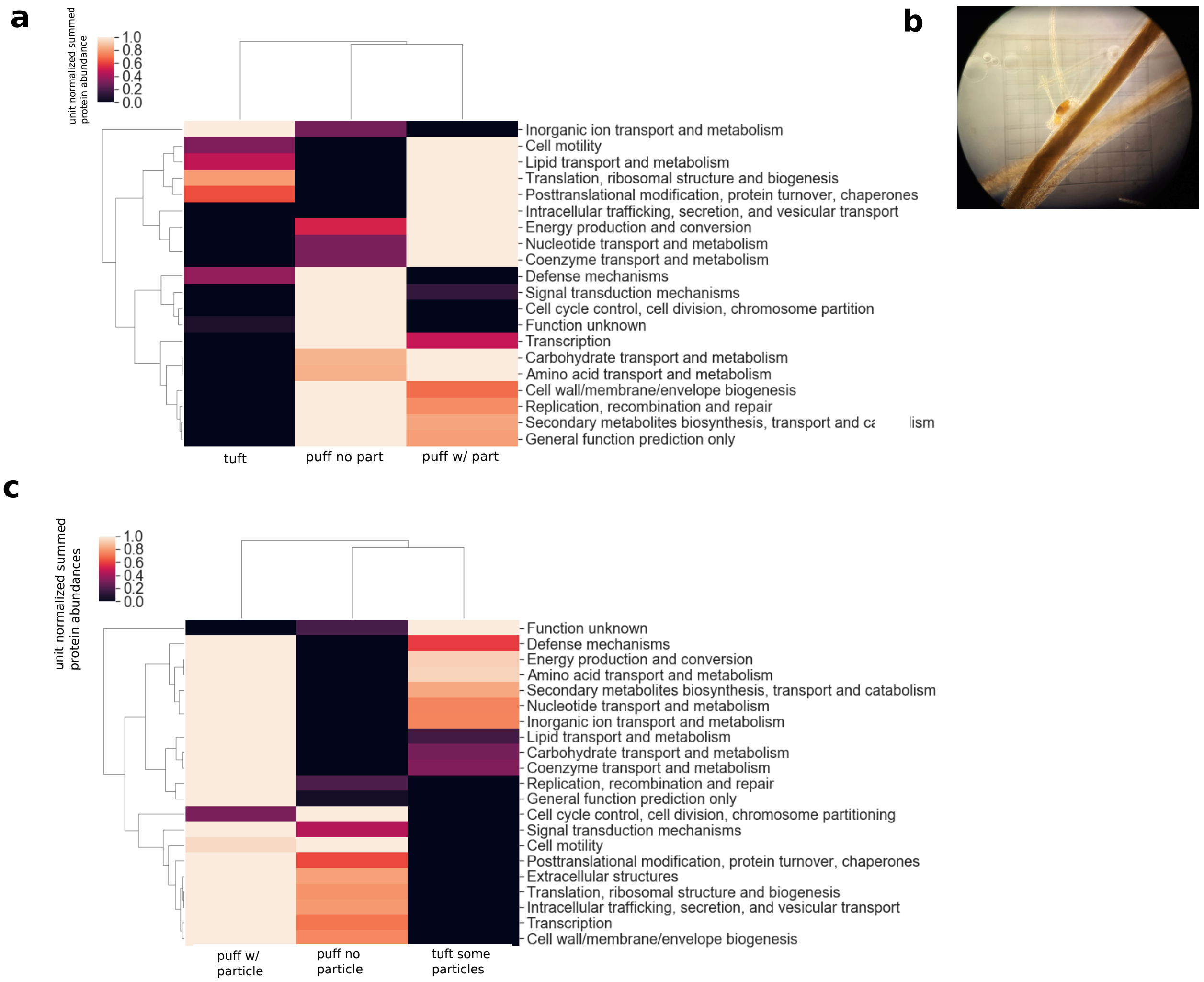


**Figure S7.** A) Clustered heatmap of summed relative protein abundances of the COG functional modules for epibiont proteins only, classified by colony morphology. Summed protein abundances have been unit normalized across the row. B) Light microscopy image of metazoan (copepod) associated with a *Trichodesmium* colony from this location. C) Clustered heat map of the abundance of COG functions for the different morphology types, but for *Trichodesmium* proteins only). Abundances are the sum of the normalized protein abundances assigned to the KO module and are unit-normalized across the row.

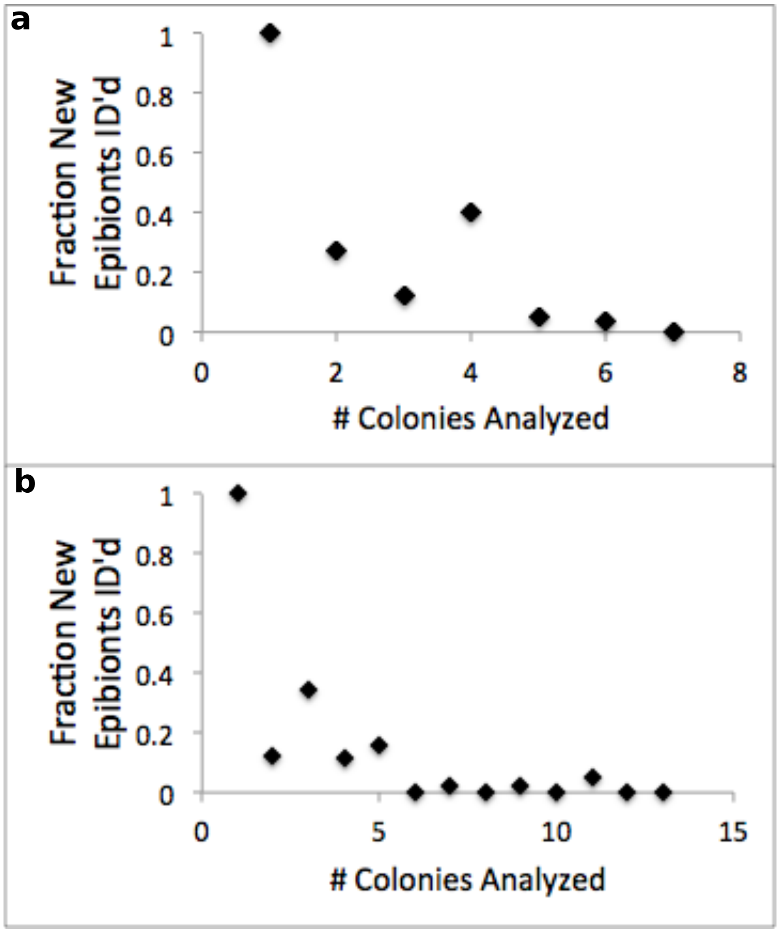


**Figure S8.** Rarefaction curve of the epibiont species identified in the individual colony metaproteomes for colonies without (top) and with (bottom) particle associations. In both cases saturation of the epibiont community was reached after just a few colonies were analyzed, indicating complete coverage of the species diversity based on the analytical workflow utilized.

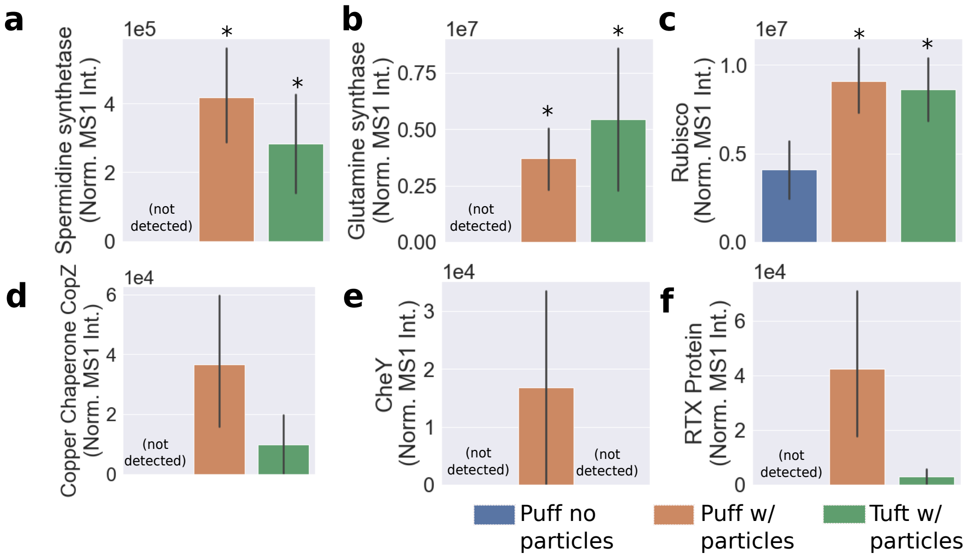


**Figure S9.** Average relative abundance of more selected proteins noted in the main or supplementary text for the different colony types. *Indicates statistically significant difference compared to the puffs without particles by a two-tailed t test, $p < 0.05$. Error bars are one standard deviation of the mean.

**Table S1.** Fe XANES fitting results for determination of Fe oxidation state. Ferrihydrite and siderite are used to represent Fe(III) and Fe(II) end-member components.

| Sample | Fe XANES Component Fits | | Average Oxidation State | $X_{\upsilon}^2$ |
|---|---|---|---|---|
| | Fe(III) Ferrihydrite | Fe(II) Siderite | | |
| Puff 1, Particle 1 | 1.00 | — | 3.0 | 0.0013 |
| Puff 1, Particle 2 | 0.87 | 0.13 | 2.9 | 0.0007 |
| Puff 1, Particle 3 | 0.57 | 0.43 | 2.6 | 0.0033 |
| Puff 2, Particle 1 | 0.91 | 0.09 | 2.9 | 0.0030 |
| Puff 2, Particle 2 | 1.00 | — | 3.0 | 0.0009 |
| Puff 2, Particle 3 | 0.69 | 0.31 | 2.7 | 0.0032 |

**Additional Supplemental Tables (provided separately)**

**Table S2.** Taxonomic distribution of metagenomic reads of *Trichodesmium* colonies from Station 19 (large file).
**Table S3.** Protein identifications and relative quantitation data for single colony metaproteomes (large file)
**Table S4.** Protein identifications and relative quantitation data for triplicate Trichodesmium population-level metaproteomes from the sampling location (large file)
**Table S5.** Last common ancestor analysis for epibiont peptides (large file).
